# Marked irregular myofiber shape is a hallmark of human skeletal muscle aging and is reversed by heavy resistance training

**DOI:** 10.1101/2023.06.05.543651

**Authors:** Casper Soendenbroe, Anders Karlsen, Rene B. Svensson, Michael Kjaer, Jesper L. Andersen, Abigail L. Mackey

## Abstract

**Background:** Age-related loss of strength is disproportionally greater than the loss of mass, suggesting maladaptations in the neuro-myo-tendinous system. Myofibers are often misshaped in aged and diseased muscle, but systematic analyses of large sample sets are lacking. Our aim was to investigate myofiber shape in relation to age, exercise, myofiber type, species, and sex.

**Methods:** Previously collected vastus lateralis muscle biopsies (n=265) from 197 males and females, covering an age-span of 20 to 97 years, were examined. The gastrocnemius and soleus muscles of 7 C57BL/6 mice were also examined. Immunofluorescence and ATPase stainings of muscle cross-sections were used to measure myofiber cross-sectional area (CSA) and perimeter, from which a shape factor index (SFI) was calculated in a fiber type specific manner (type I and II in humans; type I, IIa, IIx and IIb in mice). Heavy resistance training (RT) was performed 3 times per week for 3-4 months by a subgroup (n=59). Correlation analyses were performed comparing SFI and CSA with age, muscle mass, maximal voluntary contraction (MVC), rate of force development (RFD), and specific force (MVC/muscle mass).

**Results:** In human muscle, SFI was positively correlated with age for both type I (R^2^=0.20) and type II (R^2^=0.38) myofibers. When subjects were separated into age cohorts, SFI was lower for type I (p<0.001) and II (p<0.001) myofibers in Young (20-36) compared to Old (60-80), and higher for type I (p<0.05) and II (p<0.001) myofibers in the Oldest Old (>80) compared to Old. The increased SFI in old muscle was observed in myofibers of all sizes. Within all three age cohorts, type II myofibers SFI was higher than for type I myofibers (p<0.001), which was also the case in mice muscles (p<0.001). Across age cohorts, there was no difference between males and females in SFI for either type I (p=0.496/0.734) or II (p=0.176/0.585) myofibers. Multiple linear regression revealed that SFI, after adjusting for age and myofiber CSA, has independent explanatory power for 8 out of 10 indices of muscle mass and function. RT reduced SFI of type II myofibers in both Young and Old (p<0.001).

**Conclusions:** Here, we identify type I and II myofiber shape in humans and mice as a hallmark of muscle ageing, that independently predicts volumetric and functional assessments of muscle health. RT reverts the shape of type II myofibers, indicating that lack of neuromuscular activation might lead to myofiber deformity.

## Introduction

It has been known for at least 50 years that myofibers atrophy with ageing^1^, and today, type II myofiber atrophy is a recognized hallmark of muscle ageing^2^. In addition to atrophy, the number of myofibers irreversibly declines with age^3^, due to loss of innervation^4^. Consequently, muscle mass and function, which is critical for independent living, diminish with increasing age^5^. Sarcopenia, the clinical diagnosis for severe loss of muscle mass, strength, and function, is observed in 5-10% of individuals >65 years of age and in >25% of individuals >80 years of age^6^, which highlights the importance of preserving muscle mass and function with ageing. An interesting aspect of these age-related changes is that the loss of muscle function is 2-4 fold greater than the loss of muscle mass^5,7,8^, meaning that for a given amount of muscle, less force is produced in older compared to younger adults (i.e., lower functional muscle quality). This indicates the presence of maladaptations in the neuro-myo-tendinous system.

One potential such maladaptation is how myofibers are shaped. Geometric analyses of tissue architecture have shown that physical constraint restricts some tissues, including skeletal muscle, into specific organizations^9,10^. Myofibres are collectively organized in tessellations of convex polygons^9^. Crucially, if the physical constraint that maintains organization of packed tissues (such as skeletal muscle) is lifted, deviations from the normal shape will occur. One example of this is seen when neuromuscular transmission is lost, which initiates atrophy and ultimately loss of the myofiber^11^. Consequently, it is common in many neurogenic and myogenic genetic disorders with denervation, to observe atrophic and deformed myofibers amidst myofibers with normal morphology^12^. Furthermore, we have observed that such fibres express denervation markers and their prevalence increases with ageing in healthy cohorts^4,13,14^. Interestingly, a potential explanation for the presence of misshaped fibres is that the healthy innervated myofibers compress denervated myofibers^10^.

Four studies have investigated myofiber shape in humans^15–18^. Type II myofibers are more deformed in older compared to younger adults, while the shape of type I myofibers appears to be less affected by ageing^15–17^. Additionally, there does not seem to be differences in myofiber shape between males and females^16,17^. Shape might however be influenced by an individual’s level of physical activity^17^, although it has earlier be reported that a 9-week resistance exercise training intervention did not alter myofiber shape^18^. Furthermore, several older studies have provided qualitative descriptions of increased myofiber deformity in both rodents and humans, relating to ageing^19–21^, cachexia^1^, denervation^22,23^ and muscular dystrophies^24^. In general, for these studies, it is mostly – if not solely – the shape of type II myofibers that seems to be affected, but systematic analyses of myofiber shape in large data sets are lacking.

Currently, it is unknown how myofiber shape is related to physical function and thus whether it represents an undescribed hallmark of aging. Similarly, the number of studies investigating age-related changes in myofiber shape is low, and it is completely unknown whether hypertrophy inducing resistance training affects myofiber shape. As such, the aim of this study was to investigate how myofiber shape is influenced by ageing and fiber type in both young, old, and very old males and females, with varying levels of physical function. Additionally, hypertrophy inducing resistance training was used as a modality that might positively impact myofiber shape. We hypothesized that the shape of type I and II myofibers would be similar in young, healthy individuals, and that ageing would lead to deformation of myofibers, especially in type II myofibers. Furthermore, we expected that hypertrophy inducing resistance training could reverse the deformation of myofiber shape.

## Methods

### Ethical approval and participants

For this study, 265 muscle biopsies collected from 197 individuals that were part of 7 prior studies previously conducted at Bispebjerg and Frederiksberg hospitals^13,25–30^, were reanalyzed. All studies were approved by appropriate ethics committees (Ref: (KF) 01-224/94, H-4-2013-068, H-15005016, H-15005761, H-15017223, H-16014268, H-19000881) and were conducted in accordance with the standards set by the Declaration of Helsinki. All subjects signed an informed consent agreement prior to enrollment.

The inclusion and exclusion criteria varied between studies but in general, young^13,25,27^ and most older adults^13,25–27^ were healthy, non-smoking, medication-free, and largely physically inactive males and females aged 20-36 or >60 years. They did not have any knee or hip pain which might prohibit physical testing, use anticoagulant medication, have a high alcohol consumption, or have a BMI outside the normal range (<18.5 or >34.0 kg/m^2^). Some older adults lived in nursing homes, and had varying levels of assistance with everyday activities and food preparation^28,30^. These participants were medically examined prior to enrollment, to make sure that they did not have surgical or medical diseases that prevented participation. Finally, some of the older adults were patients admitted to the geriatric ward, who were included in the study primarily based on whether they could safely and capably complete the study^29^.

### Heavy resistance training

Young (n=7) and older (n=52) adults underwent 3-4 months of heavy resistance training (RT), three times per week^25,26^. Following 5 minutes of low to moderate intensity cycling, the participants performed 3-5 sets of 8-15 repetitions at 8-15 repetition maximum, in leg press, leg extension, leg curl and two optional upper body exercises. The training was supervised, and loads were continuously increased when a target number of repetitions at a given load was achieved.

### Muscle mass, strength, and specific force

Leg lean mass (LBM_leg_) was measured by Dual Energy X-Ray Absorptiometry (DEXA) in 108 out of 197 subjects. Cross sectional area of the quadriceps (CSA_Quad_) was by Magnetic Resonance Imaging (MRI) in 104 out of 197 subjects^25^. Maximal voluntary contraction force (MVC) was measured in 149 out of 197 subjects, using a dynamometer (Model 500−11; Kinetic Communicator) in both isokinetic (60°/s) and isometric (at 70° knee extension, 0° equals straight leg) mode^13^. Peak rate of force development (RFD) was simultaneously derived from the isometric MVC. Specific force (strength per unit mass) was calculated as isokinetic or isometric MVC relative to LBM_leg_ or CSA_Quad_.

### Muscle biopsy

Muscle biopsies were obtained from the mid-belly of the vastus lateralis muscle, under local anesthesia (lidocaine), using a 5 or 6 mm. Bergstrom needle, fitted with manual suction. The biopsied tissue was aligned in Tissue-Tek (Sakura Finetek), and frozen in isopentane (JT Baker) pre-cooled by liquid nitrogen. Samples were stored at -80°C.

### Animals

Seven 11-months old C57BL/6 mice (Janvier), that were part of a previous study (Danish Animal Inspectorate, Ministry of Justice (#2014-15-020100326)^31^, were housed individually, under standard conditions (tap water and standard chow) without assess to running wheel in a 12h light-dark cycle. The mice were sacrificed by cervical dislocation after which the left gastrocnemius (GAS) and soleus (SOL) muscles were dissected and aligned in Tissue-Tek (Sakura Finetek), and frozen in pre-cooled isopentane (JT Baker). Samples were stored at -80°C.

### Histochemistry, immunofluorescence, and imaging

Muscle samples (human biopsies and mouse muscles) were sectioned (7-10 μm thickness) cross sectionally using a cryostat, mounted on SuperFrost® Plus glass slides (Menzel-Gläser, Thermo Scientific) and stored at -80 °C. Human muscle samples were fiber typed using either ATPase histochemistry (97 samples) or immunofluorescence (168 samples).

#### ATPase – human tissue

ATPase histochemistry and visualization of the myofiber membrane was performed as previously described^26,30^. Briefly, immunohistochemical staining of the myofiber membrane and capillaries were performed using a double staining method combining ulex europaeus lectin 1 (UEA-1) and collagen type IV staining^32^. ATPase histochemistry for identifying fiber types was performed by preincubating four slides with solutions of pH 4.37, 4.53, 4.57, and 10.30, respectively^26^. Using the immunohistochemical staining for the myofiber membrane, a fiber mask was drawn along the cell borders of the desired number of fibers.

Images of the ATPase stainings were then fitted into the fiber mask. A number was assigned by the computer to each fiber and the fibers were then displayed on the screen in multiple images. The individual fibers were therefore easily identified and assigned to a specific fiber type. Myofiber cross-sectional area (CSA) and perimeter was measured using an image-analysis software (Tema, Scanbeam, Hadsund, Denmark). Type II isotypes (IIa and Iix) were pooled into type II. All type I/Iia hybrid fibers, which are common in aged muscle^33^, were removed from quantification. A mean ± SD [range] of 73 ± 24 [21-125] type I and 67 ± 23 [18-154] type II fibers per sample were analyzed by ATPase.

#### Immunofluorescence – human tissue

Immunofluorescence was performed as previously described^13,29^. Briefly, sections were removed from the freezer and allowed to dry. Dystrophin (RRID:AB_259245, D8168, Sigma-Aldrich) or laminin (RRID:AB_2313665, Z0097, Dako) was used as membrane marker together with a MyHC I antibody (RRID:AB_10540570, A4.951 or RRID:AB_2235587, BA.D5, both DSHB). Primary antibodies were appropriately diluted in a blocking buffer consisting of 1% bovine serum albumin (BSA) and 0.1% sodium azide in Tris-buffered saline (TBS) and applied overnight at 5°C. The next day, slides were incubated for 45 min at room temperature with corresponding secondary antibodies appropriately diluted in blocking buffer. Finally, sections were mounted with cover glasses using Prolong-Gold-Antifade (P36931; Thermo Fisher Scientific). Fixation was performed using either 4% PFA or Histofix (Histolab) either before primary antibodies or after secondary antibodies were applied. Sections were washed in TBS between all steps. Imaging was performed using either a 0.30 NA/10× objective and a DP71 Olympus camera on a BX51 Olympus microscope, or 0.80 NA/20x objective on a confocal laser scanning microscope (LSM710, Carl Zeiss, Oberkochen, Germany). Image analysis was performed using a custom-build ImageJ macro^34^, as previously described in detail^35^. Type I (MyHC I positive) and type II (MyHC I negative) myofiber CSA and perimeter was measured. Hybrid myofibers (weak MyHC I staining) were removed. A mean ± SD [range] of 231 ± 125 [38-710] type I and 204 ± 127 [31-705] type II myofibers per sample were analyzed by immunofluorescence.

Myofiber denervation was assessed in a subset of human samples using immunofluorescence to detect the presence of neural-cell-adhesion-molecule (NCAM) in myofibers, as previously described in detail^4^.

#### Immunofluorescence – mouse tissue

For mouse muscles, as previously described^31^, two consecutive cross sections were stained with laminin (RRID:AB_2313665, Z0097, Dako) and either MyHC1 (RRID:AB_2235587, BA.D5, DSHB) and MyHC2b (RRID:AB_2811120, BF-F3, DSHB), or MyHC2a (RRID:AB_2147165, SC-71, DSHB) and MyHC2x

(RRID:AB_1157897, 6H1, DSHB). Samples were imaged using a 0.50 NA/20× objective and a DP71 Olympus camera on a BX51 Olympus microscope. Images were analyzed using a custom-made semi-automated macro in ImageJ, which aligned images from consecutive cross sections, identified fiber type and performed morphometric measurements. A mean ± SD [range] of 29 ± 22 [2-59] type I and 1048 ± 373 [355-1438] type II myofibers per GAS sample, and 251 ± 105 [129-444] type I and 375 ± 133 [162-523] type II myofibers per SOL samples, were analyzed.

### Shape factor index

Myofiber shape was evaluated using the shape factor index (SFI) as previously performed^15–18^. SFI is a derivative of the “Formfactor”, and thus represents a dimensionless expression of object shape, with values above 1.00 showing deviation from a circle^36^. Myofibers cut in the cross-sectional plane have polyhedral shapes rather than circular, and have values above 1.00, with greater values indicating further shape deviation. An increased SFI thus means that a myofiber has a greater perimeter relative to its area. SFI was calculated for each myofiber using the measured CSA and perimeter values, with the formula:

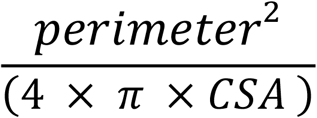

Where the denominator represents the squared perimeter of a perfect circle with the given CSA (the smallest possible perimeter). Being a dimensionless expression of shape, objects of the same shape but different sizes will have the same SFI (fig.1.A). But two myofibers of equal size can have greatly different shapes (fig.1.B). To exemplify further, fig.1.C shows the SFI of a circle (1.00), pentagon (1.15), hexagon (1.11) and myofibers with gradually increasing SFI (1.14-1.87).

**Fig. 1.**
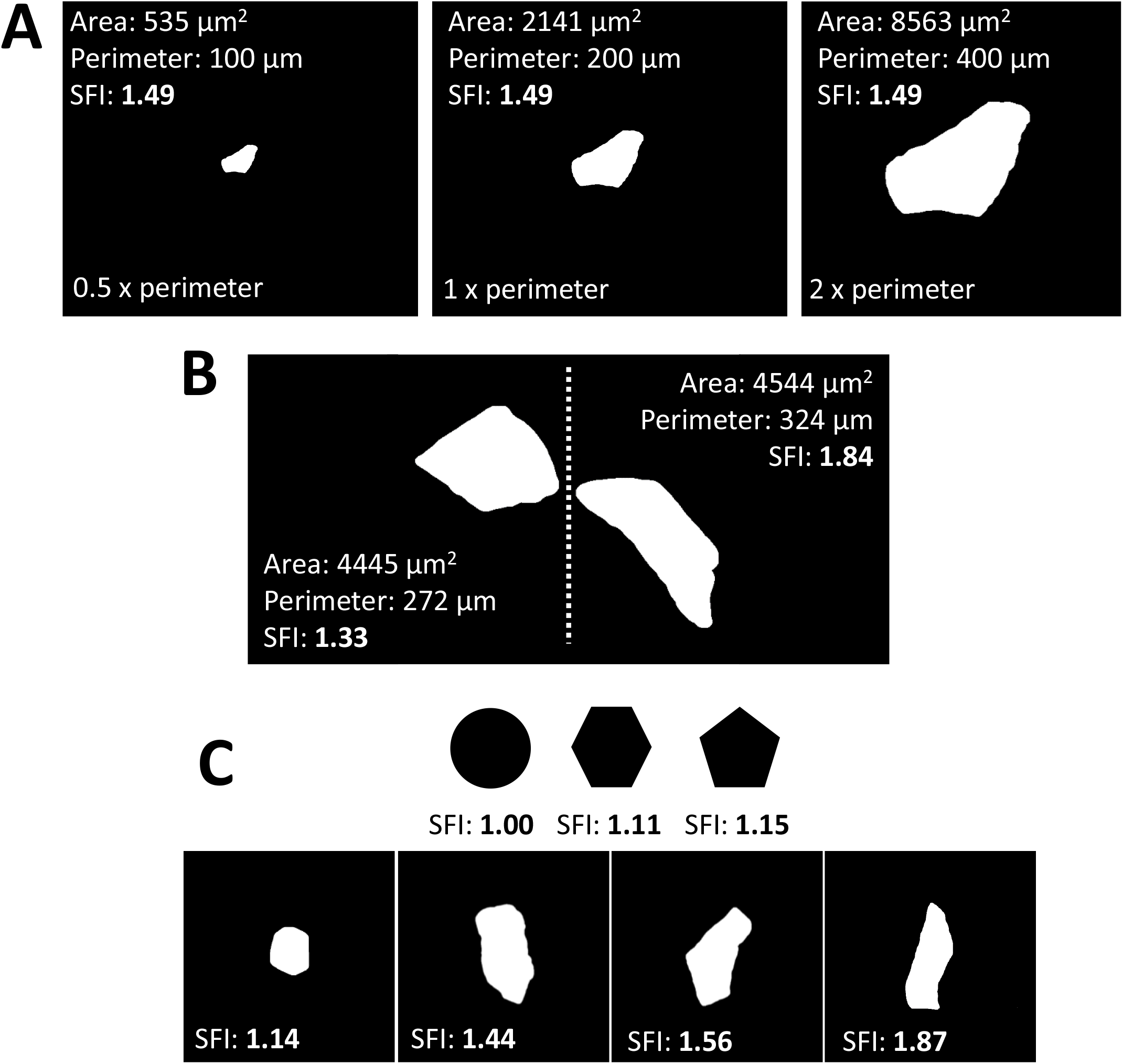
Shape Factor Index (SFI) overview. A) SFI is a dimensionless descriptor of shape, as shown with the same object in three different sizes. B) High SFI values indicate increased deviation from a circle, as illustrated by two fibers with similar CSA but different shapes. C) SFI values for a circle and four myofibers of different shapes. CSA, cross-sectional area.

For each tissue sample, the mean SFI of all myofibers analyzed was calculated to represent that sample. Furthermore, SFI distribution was expressed in 0.10 increments from <1.10 to >2.00, as well as in relation to myofiber CSA in 1000 μm^2^ increments from 0 to >6000 μm^2^.

### Statistics

Data in graphs are presented as means ± SEM or means with individual values. Data were analysed using t-tests (paired or unpaired), one-way ANOVAs, two-way repeated measures ANOVAs and mixed-effects models. The statistical test used for a given dataset is specified in the corresponding figure legend. Relationship between SFI, or CSA, and muscle mass and function was evaluated using simple regressions (Pearson), whereas the association between SFI and age was evaluated using second order polynomial regression analyses. Multiple linear regressions (backward elimination) were performed to explore the explanatory power of five model parameters (age, type I SFI, type II SFI, type I CSA and type II CSA) on muscle mass and function. Type II semi-partial correlation coefficients, which represent the additional explanatory power of the given parameter after accounting for the other parameters in the model, are reported for significant predictors. Graphs were prepared in Prism (v.8, GraphPad Software). Statistical analyses of the histograms (two-way repeated measures ANOVAs and mixed-effects models) were performed in Prism, multiple linear regressions were performed in SAS (v.9.2, SAS, Cary, NC, USA), and all other statistical analyses were performed in SigmaPlot (v. 13.0, Systat Software). Statistical significance was set at 0.05.

## Results

### Subject characteristics

In the present study, muscle biopsies from 197 subjects, that were part of 7 prior studies, were analyzed for SFI. Key subject characteristics are summarized in table 1. The number of female (F) and male (M) subjects represented in each age cohort was 12 F and 22 M in Young, 13 F and 98 M in Old, and 22 F and 30 M in Oldest Old. Sex-specific subject characteristics are presented in online supporting information 1. The Old and Oldest Old groups were phenotypically aged; compared to Young, Old demonstrated lower LBM_leg_ (11%, p<0.05), CSA_quad_ (16%, p<0.05), and isometric MVC (21%, p<0.001). Compared to Old, Oldest Old demonstrated lower LBM_leg_ (21%, p<0.001), CSA_quad_ (26%, p<0.001), and isometric MVC (33%, p<0.001). Subject characteristics from each of the 7 studies are provided separately in online supporting information 2.

**Table 1.** Subject characteristics for Young, Old, and Oldest Old shown as means ± SD with range and number of subjects represented in each measurement. LBM_leg_, CSA_quad_, MVCs and RFD were compared using one-way ANOVAs. * p<0.05 between Young and Old. # p<0.05 between Old and Oldest Old. Subject characteristics separated by gender and from each study can be found in online supporting information 1 and 2, respectively. BMI, body mass index. LBM_leg_, leg lean mass. CSA_quad_, quadriceps cross-sectional area. MVC, maximal voluntary contraction. RFD, rate of force development.

|  | Young (12F/22M) |  |  | Old (13F/98M) |  |  | Oldest old (22F/30M) |  |  |
| --- | --- | --- | --- | --- | --- | --- | --- | --- | --- |
| | Average ± SD | Min - Max | N | Average ± SD | Min - Max | N | Average ± SD | Min - Max | N |
| Age (y.) | 25 ± 4 | 20 - 36 | 34 | 71 ± 4 | 60 - 79 | 111 | 86 ± 3 | 81 - 97 | 52 |
| Height (m.) | 1,77 ± 0,10 | 1,57 - 1,93 | 34 | 1,77 ± 0,08 | 1,55 - 1,95 | 111 | 1,70 ± 0,11 | 1,42 - 1,88 | 40 |
| Weight (kg.) | 75 ± 15 | 53 - 111 | 34 | 81 ± 13 | 56 - 109 | 111 | 72 ± 14 | 45 - 98 | 40 |
| BMI (kg/m<sup>2</sup>) | 23,9 ± 3,1 | 19,0 - 30,4 | 34 | 25,7 ± 3,3 | 18,9 - 32,5 | 111 | 24,6 ± 3,3 | 18,7 - 33,2 | 40 |
| LBM<sub>leg</sub> (kg.) *# | 22 ± 4 | 17 - 30 | 22 | 20 ± 3 | 12 - 26 | 53 | 16 ± 3 | 9 - 21 | 33 |
| CSA<sub>quad</sub> (cm<sup>2</sup>) *# | 73 ± 15 | 55 - 99 | 7 | 62 ± 12 | 36 - 91 | 65 | 45 ± 11 | 26 - 69 | 32 |
| Isometric MVC (Nm) *# | 272 ± 62 | 186 - 364 | 22 | 188 ± 38 | 85 - 278 | 93 | 126 ± 42 | 22 - 205 | 34 |
| Isokinetic MVC (Nm) *# | 307 ± 67 | 200 - 425 | 22 | 207 ± 46 | 75 - 313 | 94 | 109 ± 45 | 18 - 219 | 35 |
| Isometric RFD (Nm/s) * | 2322 ± 839 | 1135 - 4048 | 22 | 1259 ± 472 | 251 - 2746 | 93 | 1126 ± 531 | 692 - 2295 | 8 |

### SFI is fiber type dependent and increases with ageing

SFI for both type I and II myofibers demonstrated positive correlations with age (fig. 2.A; R^2^ = 0.20 and 0.38, p<0.001), with myofibers becoming increasingly misshapen with age. There was no difference in SFI between males and females in Young, Old, and Oldest Old (online supporting information 3). To further explore the different trajectories between fiber types, subjects were divided into three age cohorts (fig. 2.B), where it became clear that the fiber type difference in SFI was present even in Young (p<0.001). Additionally, Old had higher SFI values for both fiber types than Young (p<0.001), while the Oldest Old had higher values than Old (p<0.05-0.001). The age-related increase in SFI was smaller for type I compared to type II myofibers. Specifically, compared to Young, type I SFI was 3.9 % and 5.2 % higher in Old and Oldest Old, respectively. For type II fibers, the equivalent values were 6.2 % and 13.6 %. Examples of fibers with the full range of SFI values are shown in figure 2.C-E and online supporting information 4.

**Fig. 2.**
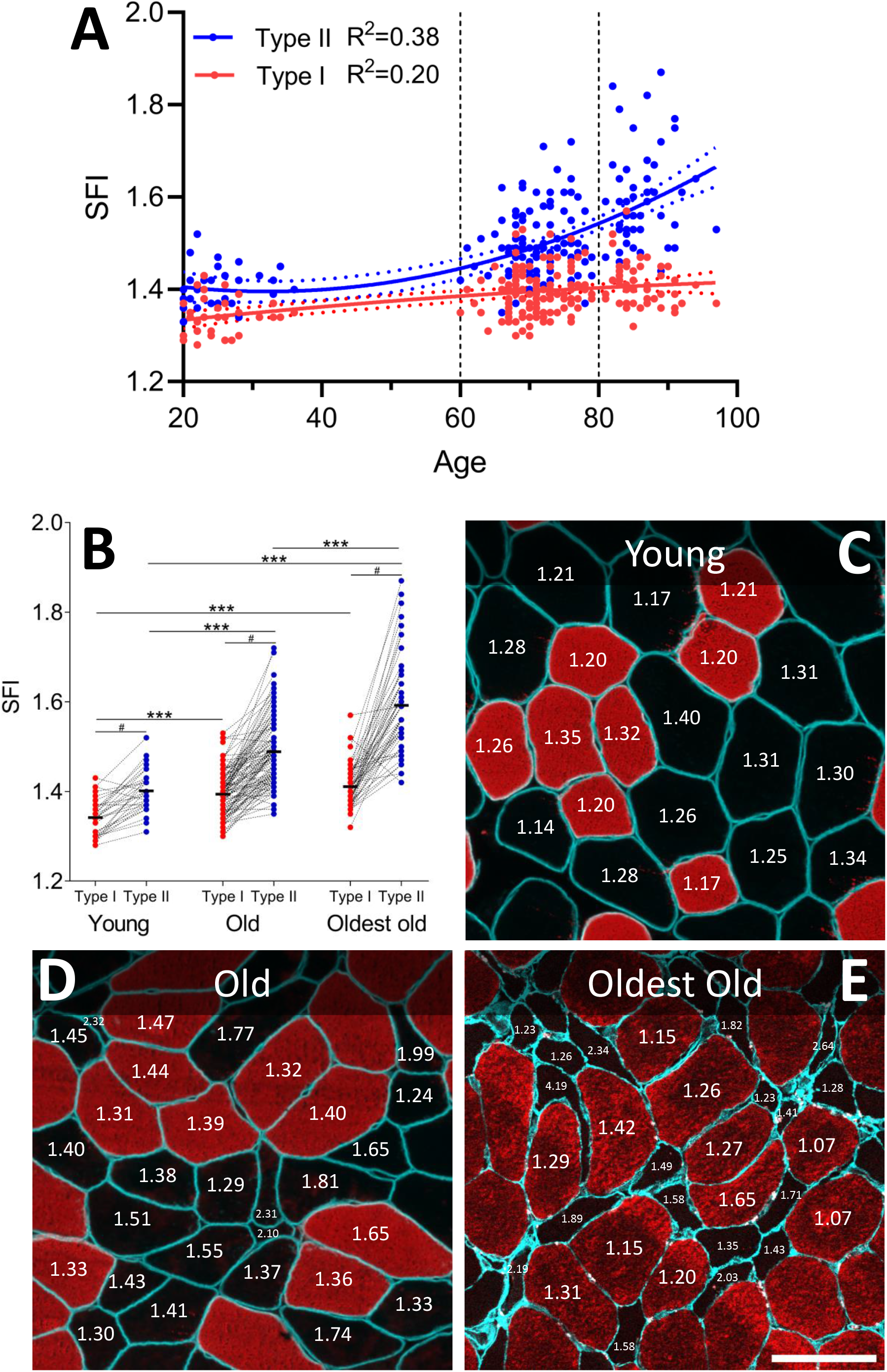
Myofiber SFI increases with ageing and in type II fibers. A) Association between age and type I (red, p<0.001) and II (blue, p<0.001) myofiber SFI was determined by second order polynomial regression (solid line). Horizontal dashed lines indicate arbitrary separation into three age cohorts: Young (n=34), Old (n=111) and Oldest Old (n=52). B) Type I and II myofiber SFI for Young, Old, and Oldest Old displayed as means, with connected individual values. Each data point represents one individual. Data were analyzed using Two Way Repeated Measures ANOVA (age group x fiber type), with significant main effects (p<0.001) and interactions (p<0.001). *** indicate effect of age group, p<0.001. # indicate effect of fiber type, p<0.001. C-E) Representative examples of muscle biopsy cross-sections from Young (C), Old (D), and Oldest Old (E), stained for dystrophin (C-D, cyan), laminin (E, cyan) and MyHC I (C-E, red). SFI values are provided for selected fibers. Scale bar = 100 μm. SFI, shape factor index. MyHC I, myosin heavy chain type I.

### SFI is higher in older individuals for fibers of all sizes

With increasing age, fibers with high SFI values make up a larger proportion of fibers (figure 3.A). For type I fibers, Young have a higher proportion than the two older groups in the SFI increments below 1.3. From the increments starting at 1.4 and higher, this pattern is reversed. A similar picture is clear for type II fibres, with the shift occurring at SFI of 1.5. Interestingly, only one of the type I fiber SFI increments demonstrates a difference between the Old and Oldest Old, while 7 of the type II fibre SFI increments display this difference. To probe whether the age-related change in SFI was driven by a higher abundance of atrophic fibers, SFI was expressed according to myofiber CSA, in 1000 μm^2^ increments. As seen in fig. 3.B, the greater SFI in Old compared to Young, and Oldest Old compared to Old, was present in the majority of CSA increments, confirming that high SFI is not driven by a higher prevalence of atrophic fibres alone. The number of fibers and subjects represented in each category is provided in online supporting information 5.

**Fig. 3.**
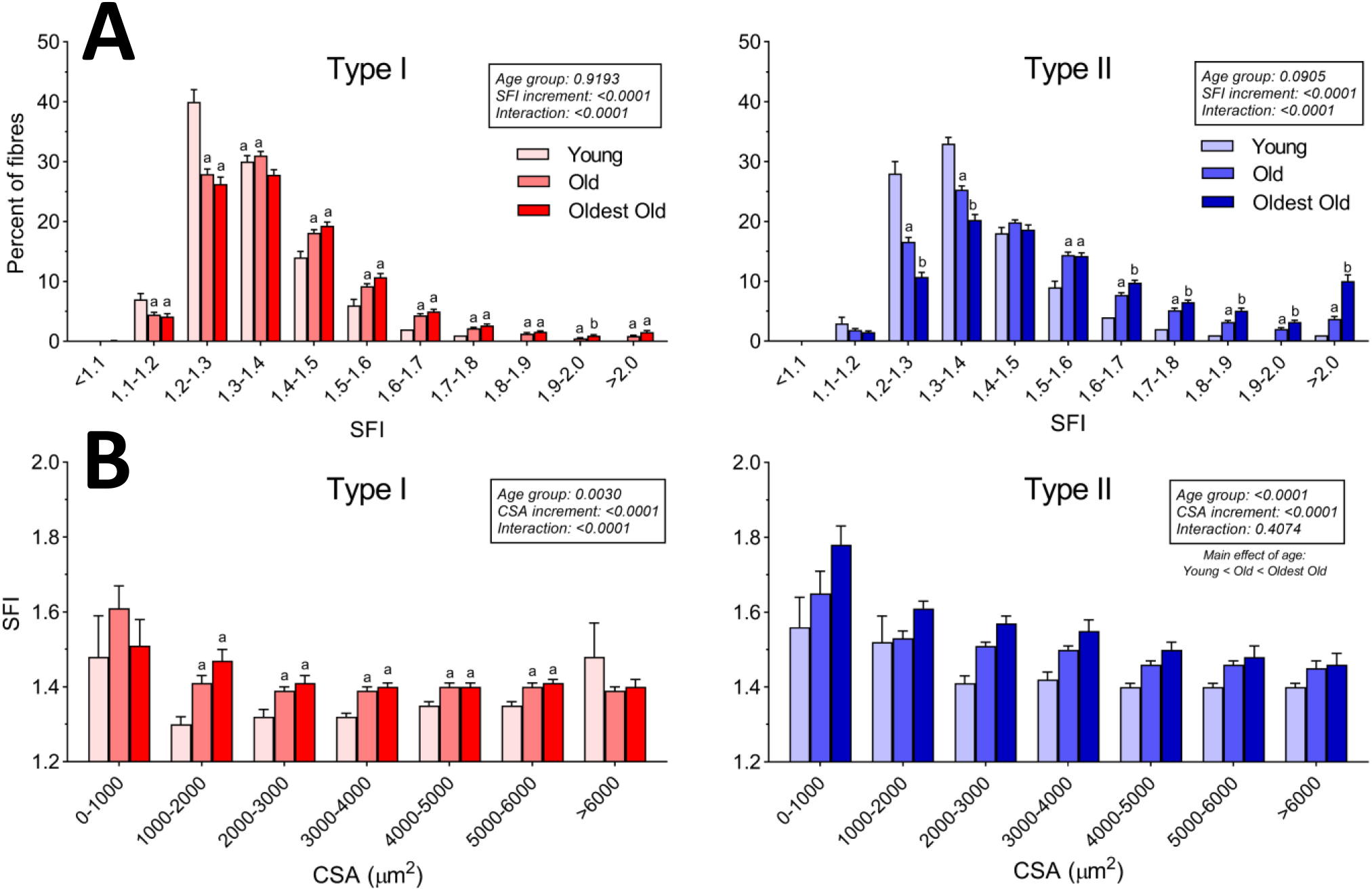
SFI distribution. A) Percentage of type I and II myofibers in 0.1 increments of SFI, for Young (n=34), Old (n=111) and Oldest Old (n=52). Data were analyzed using Two Way Repeated Measures ANOVA (age group x CSA increment), with main effects and interactions indicated in the figure. Results of the posthoc test are indicated by letters; bars that do not have the same letter are significantly different within the respective SFI increment (p<0.05). B) SFI of type I and II myofibers binned in 1000 μm^2^ CSA increments. Data are averages of all subjects within each age group and presented as means ± SEM. Data were analyzed using mixed-effects model (age group x CSA increment), with main effects and interactions indicated in the figure. Results of the posthoc test are indicated by letters; bars that do not have the same letter are significantly different within the respective CSA increment (p<0.05). See supporting information 5 for details on the numbers of participants represented in each increment. SFI, shape factor index. CSA, cross-sectional area.

We also assessed SFI for denervated myofibers (determined by NCAM expression) in a subset of samples (online supporting information 6). Most denervated myofibers are atrophic and present with high SFI values (>1.60), while a smaller portion of myofibers had a shape corresponding to that found in Young (SFI 1.20-1.50).

### Myofiber SFI shows similar patterns in mice

To address whether this was conserved across species we sampled GAS and SOL muscles from C57BL/6 mice (online supporting information 7). In confirmation of the observations in humans, SFI values were higher in type II compared to type I myofibers (p<0.001). Additionally, in the fast-twitch GAS, this seemed to vary with type II isotypes, with increasing values from IIa to IIx and IIb (p<0.05).

### The importance of SFI for physical function

To better understand how an increased SFI influences muscle function in humans, SFI and CSA were correlated with *in vivo* measurements of muscle mass (DEXA and MRI), muscle function (MVC and RFD), and specific force (strength per unit mass). As shown in figure 4, and in online supporting information 8, in general, SFI demonstrated strong, significant negative correlations with *in vivo* muscle mass and function measures, with higher SFI values associated with lower mass and function. CSA demonstrated strong, significant positive correlations with the same outcomes, with lower CSA values associated with lower mass and function. Multiple linear regression with backward elimination was used to probe whether SFI and CSA had independent explanatory power when adjusting for confounding variables, including age. As shown in figure 4.E and in online supporting information 9, SFI was an independent predictor of eight of the ten variables tested. Specifically, type II fiber SFI (and not type I fiber SFI) was an independent predictor of CSA_quad_, isometric MVC, isokinetic MVC and three types of specific force. Type I fiber SFI was an independent predictor of LBM_leg_, and SFI for both type I and II fibers were independent predictors of one measure of specific force. For comparison, type II fiber CSA only came out as an independent predictor of four out of the ten variables tested.

**Fig. 4.**
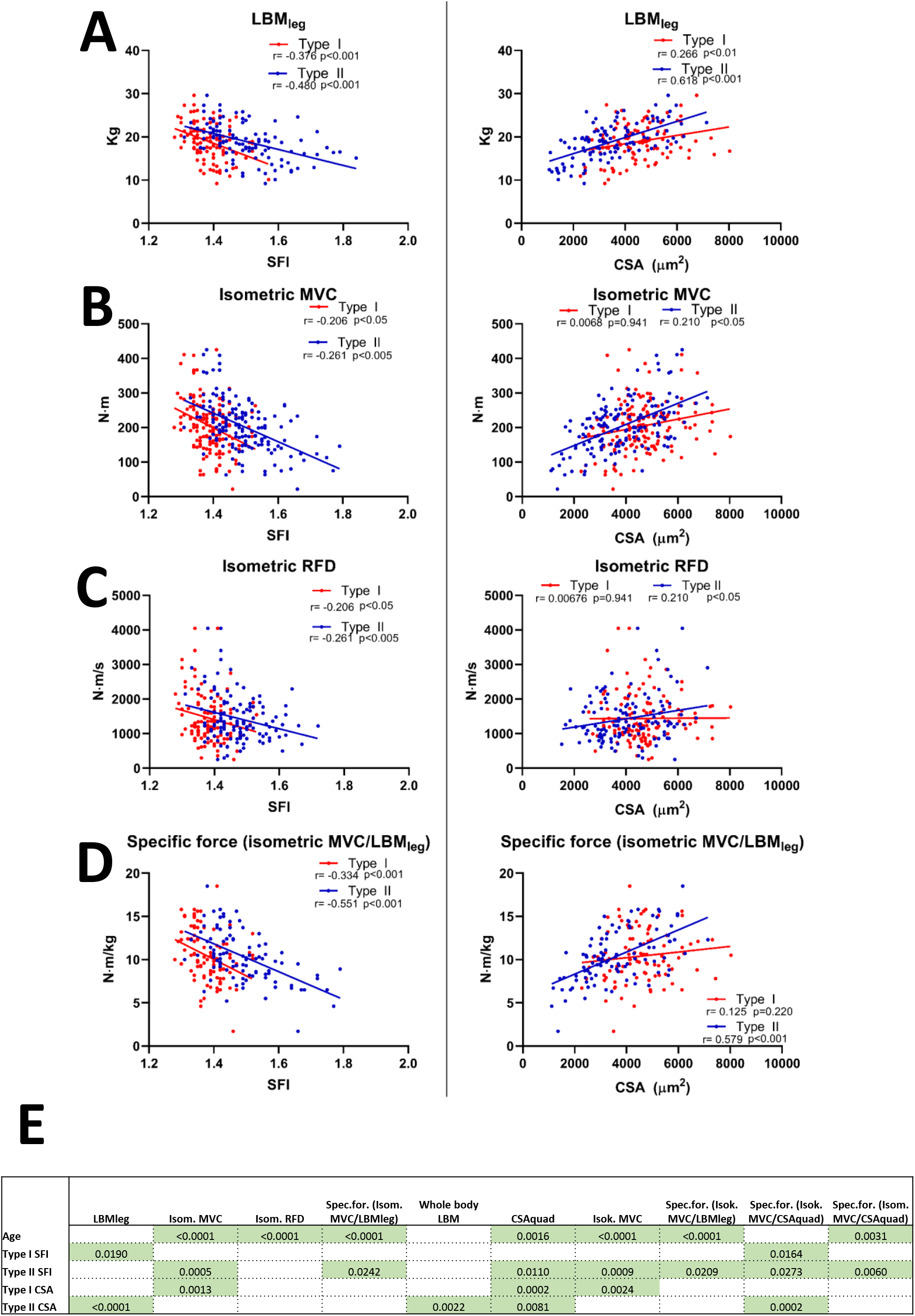
Linear correlation analyses between SFI (left) and CSA (right) for type I and II myofibers with *in vivo* measures of muscle mass and function. Each data point represents one individual, ranging in age from 20– 94y, and all in the untrained state. Strength of association is indicated by r and p values. Correlations are displayed for A) LBM_leg_ (n=108), B) isometric MVC (n=150), C) isometric RFD (n=123), D) specific force (MVC/LBM_leg_) (n=98). E) P-values of the model parameters that were significant independent predictors of ten indices of muscle mass and function. Refer to supporting information 9 for full outline of the multiple linear regression analyses. Additional correlations can be found in supporting information 8. LBM_leg_, leg lean mass. SFI, shape factor index. CSA, cross-sectional area. RFD, rate of force development. MVC, maximal voluntary contraction. Isom., isometric. Isok., isokinetic. Spec.for., specific force.

### Heavy resistance training reverses the higher SFI of type II myofibers

To test whether RT induced hypertrophy could alter myofiber shape, SFI was evaluated before and after 3-4 months of RT. In both Young and Old, RT reduced type II SFI significantly (p<0.001), while type I SFI remained unchanged (fig.5.A). Individual delta values are shown in online supporting information 10. As shown in figure 5.B, there were significant negative correlations between baseline SFI and change in SFI with RT, for both type I (R^2^=0.266, p<0.001) and II (R^2^=0.233, p<0.001) myofibers.

**Fig. 5.**
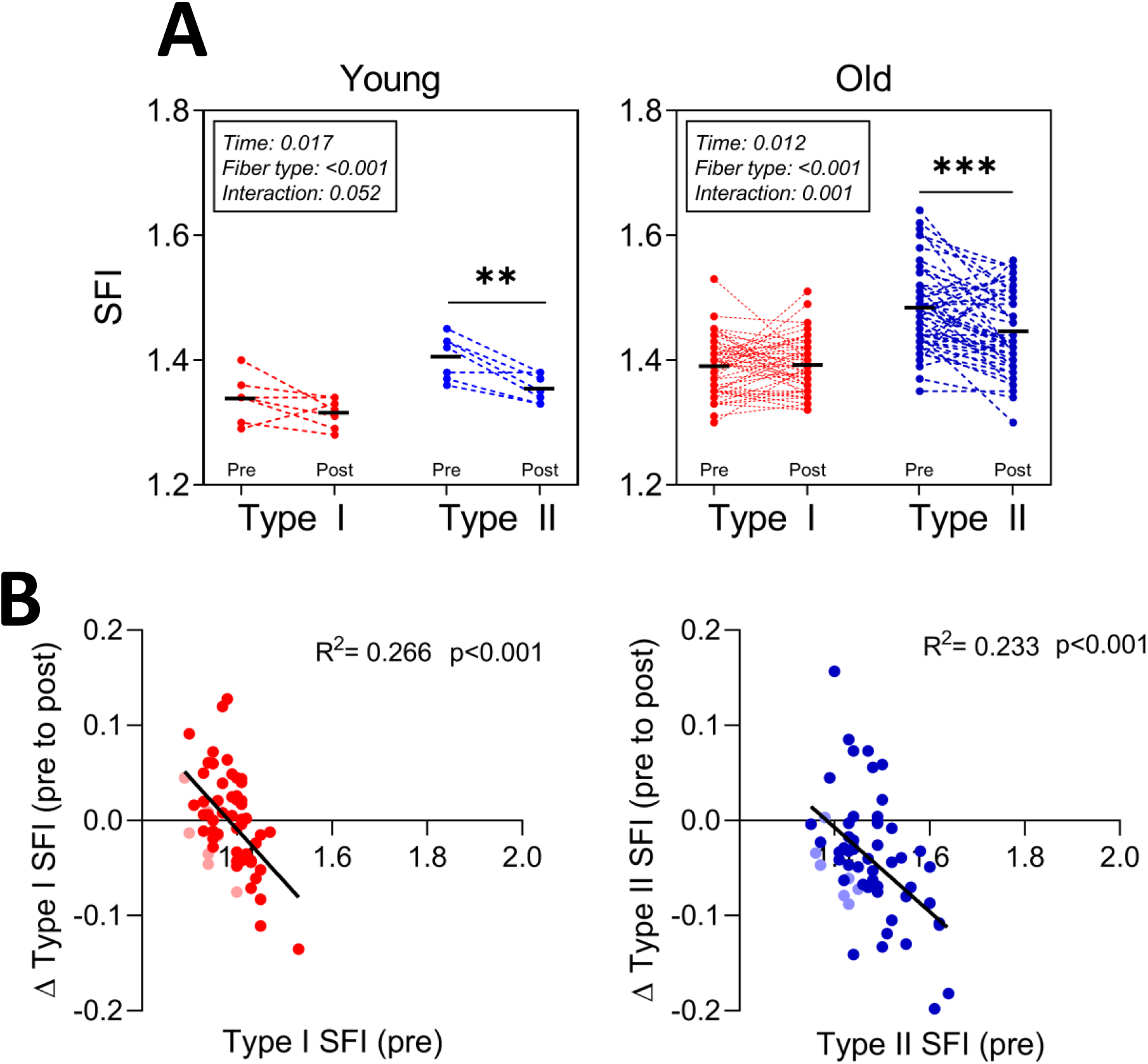
SFI modification with RT. A) SFI for type I (red) and II (blue) myofibers before and after 3 or 4 months of RT. Data are presented for Young (n=7) and Old (n=52) and were analyzed using Two-Way Repeated Measures ANOVAs (time x fiber type) within each age group, with main effects and interactions indicated in the figure. ** p<0.01 post vs pre, *** p<0.001 post vs pre. B) Linear correlation analysis between SFI at baseline and changes following RT for type I (red) and II (blue) myofibers. Data are pooled Young and Old participants from A, and Young is shown in faded colors. R^2^ and p-values are provided. RT, heavy resistance training. SFI, shape factor index.

## Discussion

Myofiber size is traditionally reported as the most important morphological feature of skeletal muscle biopsy specimens collected from healthy and diseased individuals of all ages and in response to physical activity and disuse. In this study, with 265 human muscle tissue samples, we focus instead on the shape of the myofiber and find that age, fiber type and exercise alter myofiber shape. Furthermore, myofiber shape, investigated using the shape factor index, was a significant independent predictor of several *in vivo* indices of muscle mass and function, even after adjusting for age and CSA. Importantly, hypertrophy-inducing RT partially reversed the deformity of type II myofibers. These findings demonstrate unequivocally that myofiber shape is a hallmark of muscle ageing. We argue that SFI should therefore be included alongside CSA in histological investigations of aged and diseased muscle, to enhance understanding of the full extent of myofiber maladaptation and the degree of its reversibility with mechanical loading.

### Age-related myofiber deformation is linked to lack of neuromuscular activation

It is well documented that particularly type II myofibers undergo atrophy with ageing and physical inactivity^2^, but the potential significance of myofiber shape in aged muscle has received limited attention^15–18^. In this study we comprehensively investigated myofiber shape in relation to aging, physical function, species, and sex. In contrast to our hypothesis, and previous findings^15–17^, we found that type II myofibers were deformed compared to type I myofibers, even in young healthy individuals. However, in agreement with our hypothesis but in contrast to previous findings^15–17^, we found that both type I and II myofibers became increasingly misshaped with ageing. The age-related increase in SFI was seen across all fiber size increments, albeit most predominantly in the smallest type II fibers, which are more abundant in old muscle. Age-related changes in myofiber shape are thus cumulatively driven by a general deformation of all myofibers alongside an increased proportion of small type II fibers. The age-related change in shape was less in type I fibers compared to type II. In fact, the SFI of type I myofibers in aged muscle is comparable to the SFI of type II myofibers in young. For both type I and II SFI there was a strong correlation to *in vivo* volumetric and functional assessments of muscle health. In many cases the correlations were as strong or stronger than those of the classic hallmark of muscle ageing, myofiber CSA. Consequently, it can be argued that myofiber shape is indicative of muscle health in an ageing context. Additionally, these data clearly demonstrate why it is not appropriate to use minimum Feret diameter in ageing muscle tissue samples, as the age-related change in myofiber shape will give artificially low diameters.

The increased abundance of highly deformed, and often very atrophic fibers was observed predominantly in the Oldest Old and is particularly interesting. We speculate that the source of these fibers is myofiber denervation, and failed reinnervation. When a myofiber becomes denervated, through motoneuron decay or NMJ destabilization, its gene and protein expression is altered^37^, fiber type uniformity is lost^38^, and the fiber atrophies^39^ and loses core structural features^40^. It has been shown that temporary loss of neuromuscular transmission exerts similar negative effects on myofiber size and structure as actual denervation^41^. Given that recruitment of fast-twitch motor units only occurs during high speed and high force movements, it is therefore a reasonable assumption that older adults not engaging in such activities will experience denervation-like ramifications to their type II myofibers. To explore whether the increased SFI observed in older adults was related to neuromuscular disturbances, we assessed SFI in denervated myofibers by immunofluorescence^14^. Whilst highly variable, a large proportion of the denervated myofibers were both atrophic and deformed (SFI >1.60 values). Some NCAM^+^ fibers retained a normal shape and size and are thus caught early in the denervation process. We therefore believe that the very deformed and atrophic myofibers observed are denervated and no longer able to become reinnervated.

### Inherent fiber type difference in myofiber shape

Surprisingly, we found a significant, albeit minor, fiber type difference in SFI in the young individuals, potentially due to the large number of myofibers included in the analyses (>200 per fiber type). The SFI increment with the highest proportion of type I fibers was 1.20-1.30, whilst for type II it was 1.30-1.40. This finding is surprising, as the young and healthy would be expected to have an optimal myofiber shape. We decided to investigate this further in mice and found that this inherent fiber type difference was conserved across species. Additionally, we also observed that type IIx and IIb fibers had a higher SFI than IIa fibers. It is unclear whether this reflects the divergent physiology of type I and II fiber types, with SFI relating to a fiber’s intrinsic force-generating capacity^42,43^. Alternatively, it may be acquired due to a predominately sedentary lifestyle, where fast-twitch motor units would only rarely be recruited, and this lack of neuromuscular activity could negatively influence the shape of the myofibers. Importantly, type II myofiber SFI was reduced in the 7 young subjects that underwent RT, implying that new-found recruitment and subsequent hypertrophy of these myofibers molds the myofiber from a concave to a convex polygon.

### Heavy resistance training reverses myofiber deformation

Given that SFI of both fiber types was so closely correlated with indices of physical function we wanted to explore whether hypertrophy inducing RT would normalize myofiber shape. To do that we examined muscle biopsies from young and old individuals who had completed 3 or 4 months of RT that lead to significant hypertrophy at both the whole muscle and myofiber level^25,26^. In agreement with our hypothesis, we found that SFI declined in type II myofibers following the intervention, demonstrating that increased neuromuscular activity not only results in increased size of the myofibers, but it also restores the optimal shape. It is also clear that the individuals with the highest SFI values had the largest decrements in SFI. This suggests that myofiber shape remains amendable to change even when we reach our 70s. As we did not have any subjects from the Oldest Old taking part in the training intervention, we do not know if this muscle plasticity stretches into the last decades of life.

### Conclusions and perspectives

We provide evidence in male and female humans, and in mice, to support myofiber shape – investigated using the shape factor index – as a prominent hallmark of skeletal muscle ageing. Furthermore, myofiber shape is a strong, independent predictor of indices of muscle health measured *in vivo*, and the degree of myofiber deformity is improved by several months of RT. We argue that increased myofiber deformity is related to neuromuscular disturbances and myofiber denervation. Due to the ease of assessing SFI (when muscle biopsies are collected), the strong relationship between SFI and several indices of muscle health, and the substantial adaptability of SFI in response to increased neuromuscular activity, we believe that SFI should be assessed in conjunction with other morphological muscle parameters, such as CSA. This is especially warranted when mapping the effects of interventions that seek to improve muscle quality in elderly or diseased muscle.

## Acknowledgements

Funding from the Lundbeck Foundation (R344-2020-254, R402-2022-1387) and the Nordea Foundation (Centre for Healthy Aging) is gratefully acknowledged.

The authors are thankful for the technical assistance that was provided by lab technicians Camilla Brink Sørensen, Ann-Christina Ronnie Reimann, and Anja Jokipii-Utzon provided.

The monoclonal antibodies A4.951 and BA.D5 (type I MyHC), developed by Blau, H.M., and Schiaffino S., respectively, was obtained from the Developmental Studies Hybridoma Bank, created by the NICHD of the NIH and maintained at The University of Iowa, Department of Biology, Iowa City, IA 52242.

## Author Contributions

All authors conceived and designed research; CS, AK, JLA and ALM performed experiments; CS analyzed data; CS, JLA and ALM interpreted results of experiments; CS prepared figures; CS drafted manuscript; CS, JLA and ALM edited and revised manuscript; All authors approved the final version of the manuscript.

## Conflict of interest

No conflicts of interest, financial or otherwise, are declared by the authors.

## Online supporting information

**Supporting information 1**

Subject characteristics for males and females separately, shown as means ± SD with range and number of subjects represented in each measurement. BMI, body mass index. LBM_leg_, leg lean mass, CSA_quad_, quadriceps cross-sectional area. MVC, maximal voluntary contraction.

**Supporting information 2**

Subject characteristics for each of the original studies separately, shown as means ± SD with minimum and maximum values. These subjects characteristics have previously been reported in the original publications^13,25–30^. BMI, body mass index. LBM_leg_, leg lean mass, CSA_quad_, quadriceps cross-sectional area. MVC, maximal voluntary contraction.

**Supporting information 3**

SFI in males and females separately. Type I (red) and II (blue) myofiber SFI for Young (A, n=22 males and 12 females), Old (B, n=98 males and 13 females), and Oldest Old (C, n=30 males and 22 females). Data were analyzed using Two-Way Repeated Measures ANOVA (sex x fiber type) within each group, with main effects of fiber type indicated in the figure. SFI, shape factor index.

**Supporting information 4**

Representative split channel images of muscle biopsy cross-sections from Young, Old, and Oldest Old, stained for dystrophin (Young, Old) or laminin (Oldest Old), and MyHC I. Scale bar = 100 μm.

**Supporting information 5**

A) Number of subjects and myofibers represented in each type I and II myofiber SFI increment for Young, Old, and Oldest Old. These data supplement fig. 3.A. B) Number of subjects and myofibers represented in each type I and II myofiber CSA increment for Young, Old, and Oldest Old. These data supplement fig. 3.B.

**Supporting information 6**

Split channel view of myofibers in healthy older adults stained with dystrophin (cyan), NCAM (magenta) and nuclei (white). Denervated myofiber is identified by arrow, with the SFI provided below. Scale bar = 100 μm. NCAM, neural cell adhesion molecule. SFI, shape factor index.

**Supporting information 7**

SFI in muscles of 11-month old mice. A) Type I and type II myofiber SFI in gastrocnemius (GAS) and soleus (SOL). B) SFI in GAS, for type I, IIa, IIx and IIb. C) SFI in SOL, for type I, IIa, IIx and IIb. Note that only 1 mouse had any type IIx and IIb fibers in SOL. Data are shown as means ± SEM and were analyzed using a Two-Way Repeated Measures ANOVA (muscle x fiber type), with a significant main effect of fiber type (p<0.001). * p<0.05. N=7. SFI, shape factor index.

**Supporting information 8**

Linear correlation analyses between SFI (left) and CSA (right) for type I and II myofibers with *in vivo* measures of muscle mass and function. Each data point represents one individual, ranging in age from 20– 94y, and all in the untrained state. Strength of association is indicated by r and p values. Correlations are displayed for whole body LBM, CSA_Quad_, isokinetic MVC, specific force (isokinetic MVC/CSA_Quad_), specific force (isokinetic MVC/LBM_leg_), and specific force (isometric MVC/CSA_Quad_). LBM, lean body mass. CSA_Quad_, quadriceps cross-sectional area. LBM_leg_, leg lean mass. SFI, shape factor index. CSA, cross-sectional area. RFD, rate of force development. MVC, maximal voluntary contraction.

**Supporting information 9**

Result of backward elimination multiple linear regression. Ten indices of muscle mass and function were evaluated (1-4 in fig. 4 and 5-10 in supporting information 8). Model parameters included: Age, type I SFI, type II SFI, type I CSA and type II CSA. SE, standard error. SFI, shape factor index. CSA, cross-sectional area. R squared for the individual parameters represent type II semi-partial correlation coefficients.

**Supporting information 10**

SFI delta values from pre to post RT, for type I (red) and II (blue) myofibers (n=59). Young and Old subjects are pooled, but Young is shown in faded colors. Delta values were compared using paired t-tests. *** p<0.001. RT, heavy resistance training. SFI, shape factor index.

## Notes

### Competing Interest Statement

The authors have declared no competing interest.

