## Supplemental material for "Marked irregular myofiber shape is a hallmark of human skeletal muscle aging and is reversed by heavy resistance training"

# SM 1

| | Young women (12) | | | Old women (13) | | | Oldest Old women (22) | | |
|---|---|---|---|---|---|---|---|---|---|
| | Average ± SD | Min - Max | n | Average ± SD | Min - Max | n | Average ± SD | Min - Max | n |
| Age (y.) | 23 ± 3 | 20 - 28 | 12 | 74 ± 3 | 71 - 78 | 13 | 87 ± 3 | 83 - 97 | 22 |
| Height (m.) | 1.68 ± 0.07 | 1.57 - 1.77 | 12 | 1.67 ± 0.03 | 1.62 - 1.74 | 13 | 1.59 ± 0.08 | 1.42 - 1.76 | 13 |
| Weight (kg.) | 64 ± 8 | 53 - 75 | 12 | 67 ± 9 | 57 - 84 | 13 | 57 ± 10 | 45 - 76 | 13 |
| BMI (kg/m$^2$) | 22.7 ± 2.0 | 19.0 - 25.5 | 12 | 24.2 ± 3.8 | 19.9 - 30.4 | 13 | 22.3 ± 2.9 | 18.7 - 26.2 | 13 |
| $LBM_{leg}$ (kg.) | | N/A | | 13 ± 1 | 12 - 13 | 2 | 12 ± 2 | 9 - 17 | 12 |
| $CSA_{quad}$ (cm$^2$) | | N/A | | | N/A | | 35 ± 8 | 26 - 48 | 10 |
| Isometric MVC (Nm) | | N/A | | | N/A | | 80 ± 31 | 22 - 140 | 10 |
| Isokinetic MVC (Nm) | | N/A | | | N/A | | 58 ± 19 | 18 - 91 | 10 |

| | Young men (22) | | | Old men (98) | | | Oldest Old men (30) | | |
|---|---|---|---|---|---|---|---|---|---|
| | Average ± SD | Min - Max | n | Average ± SD | Min - Max | n | Average ± SD | Min - Max | n |
| Age (y.) | 26 ± 5 | 20 - 36 | 22 | 71 ± 4 | 60 - 79 | 98 | 86 ± 3 | 81 - 93 | 30 |
| Height (m.) | 1.81 ± 0.08 | 1.68 - 1.93 | 22 | 1.78 ± 0.07 | 1.55 - 1.95 | 98 | 1.76 ± 0.07 | 1.60 - 1.88 | 27 |
| Weight (kg.) | 82 ± 15 | 62 - 111 | 22 | 82 ± 12 | 56 - 109 | 98 | 79 ± 9 | 65 - 98 | 27 |
| BMI (kg/m$^2$) | 24.6 ± 3.3 | 20.1 - 30.4 | 22 | 25.9 ± 3.2 | 18.9 - 32.5 | 98 | 25.6 ± 3.0 | 20.7 - 33.2 | 27 |
| $LBM_{leg}$ (kg.) | 22 ± 4 | 17 - 30 | 22 | 20 ± 3 | 13 - 26 | 51 | 18 ± 2 | 15 - 21 | 21 |
| $CSA_{quad}$ (cm$^2$) | 73 ± 15 | 55 - 99 | 7 | 62 ± 12 | 36 - 91 | 65 | 50 ± 8 | 37 - 69 | 22 |
| Isometric MVC (Nm) | 272 ± 62 | 186 - 364 | 22 | 188 ± 38 | 85 - 278 | 93 | 145 ± 30 | 75 - 205 | 24 |
| Isokinetic MVC (Nm) | 307 ± 67 | 200 - 425 | 22 | 207 ± 46 | 75 - 313 | 94 | 130 ± 34 | 40 - 219 | 25 |

# SM 2

| | Soendenbroe et al., 2022 (n=46) | | | |
|---|---|---|---|---|
| | Old (n=31) | | Young (n=15) | |
| | Average ± SD | Min - Max | Average ± SD | Min - Max |
| Age (yrs) | 73 ± 4 | 68 - 82 | 26 ± 5 | 20 - 36 |
| Height (m) | 1.77 ± 0.07 | 1.61 - 1.95 | 1.83 ± 0.07 | 1.69 - 1.93 |
| Weight (kg) | 79 ± 10 | 63 - 109 | 82 ± 13 | 62 - 105 |
| BMI (kg/m^2) | 25.1 ± 3.3 | 20.5 - 32.4 | 24.5 ± 3.0 | 20.3 - 29.7 |
| $LBM_{leg}$ (kg) | 20 ± 2 | 17 - 26 | 23 ± 3 | 18 - 27 |
| Isometric MVC (Nm) | 186 ± 38 | 118 - 278 | 285 ± 66 | 186 - 364 |
| Isokinetic MVC (Nm) | 197 ± 45 | 123 - 313 | 318 ± 74 | 200 - 425 |

| | Karlsen et al., 2020a (n=26) | | | |
|---|---|---|---|---|
| | Old (n=19) | | Young (n=7) | |
| | Average ± SD | Min - Max | Average ± SD | Min - Max |
| Age (yrs) | 67 ± 4 | 60 - 73 | 25 ± 3 | 20 - 31 |
| Height (m) | 1.81 ± 0.07 | 1.68 - 1.94 | 1.79 ± 0.08 | 1.68 - 1.91 |
| Weight (kg) | 85 ± 14 | 60 - 108 | 81 ± 19 | 62 - 111 |
| BMI ($kg/m^2$) | 25.9 ± 3.6 | 18.9 - 32.5 | 25.0 ± 4.2 | 20.1 - 30.4 |
| $LBM_{leg}$ (kg) | 20 ± 3 | 15 - 26 | 21 ± 5 | 17 - 30 |
| $CSA_{quad}$ ($cm^2$) | 58 ± 11 | 36 - 82 | 73 ± 15 | 55 - 99 |
| Isometric MVC (Nm) | 200 ± 36 | 130 - 257 | 244 ± 45 | 203 - 334 |
| Isokinetic MVC (Nm) | 230 ± 42 | 143 - 295 | 284 ± 44 | 230 - 358 |

| | Heisterberg et al., 2018 (n=52) | |
|---|---|---|
| | Average ± SD | Min - Max |
| Age (yrs) | 72 ± 5 | 65 - 85 |
| Height (m) | 1.78 ± 0.07 | 1.61 - 1.90 |
| Weight (kg) | 84 ± 11 | 57 - 108 |
| BMI ($kg/m^2$) | 26.5 ± 3.0 | 19.2 - 33.2 |
| $CSA_{quad}$ ($cm^2$) | 62 ± 11 | 37 - 91 |
| Isometric MVC (Nm) | 177 ± 38 | 85 - 256 |
| Isokinetic MVC (Nm) | 197 ± 47 | 75 - 298 |

| | Bechshøft et al., 2019 (n=23) | | | |
|---|---|---|---|---|
| | Old (n=11) | | Young (n=12) | |
| | Average ± SD | Min - Max | Average ± SD | Min - Max |
| Age (yrs) | 74 ± 3 | 71 - 78 | 23 ± 3 | 20 - 28 |
| Height (m) | 1.66 ± 0.03 | 1.62 - 1.69 | 1.68 ± 0.07 | 1.57 - 1.77 |
| Weight (kg) | 69 ± 10 | 57 - 84 | 64 ± 8 | 53 - 75 |
| BMI ($kg/m^2$) | 25.0 ± 3.6 | 20.3 - 30.4 | 22.7 ± 2.0 | 19.0 - 25.5 |

| | Bechshøft et al., 2017 (n=29) | |
|---|---|---|
| | Average ± SD | Min - Max |
| Age (yrs) | 87 ± 3 | 83 - 94 |
| Height (m) | 1.69 ± 0.11 | 1.42 - 1.88 |
| Weight (kg) | 71 ± 13 | 45 - 98 |
| BMI ($kg/m^2$) | 24.7 ± 3.1 | 19.0 - 30.8 |
| $LBM_{leg}$ (kg) | 16 ± 3 | 9 - 21 |
| $CSA_{quad}$ ($cm^2$) | 43 ± 10 | 26 - 69 |
| Isometric MVC (Nm) | 120 ± 45 | 22 - 205 |
| Isokinetic MVC (Nm) | 94 ± 37 | 18 - 148 |

| | Karlsen et al., 2020b (n=9) | |
|---|---|---|
| | Average ± SD | Min - Max |
| Age (yrs) | 77 ± 7 | 69 - 91 |
| Height (m) | 1.70 ± 0.09 | 1.55 - 1.83 |
| Weight (kg) | 64 ± 11 | 48 - 85 |
| BMI ($kg/m^2$) | 22.2 ± 3.2 | 18.7 - 29.1 |
| $LBM_{leg}$ (kg) | 13 ± 2 | 10 - 15 |

| | Kryger & Andersen, 2007 (n=12) | |
|---|---|---|
| | Average ± SD | Min - Max |
| Age (yrs) | 88 ± 3 | 85 - 97 |

# SM 3

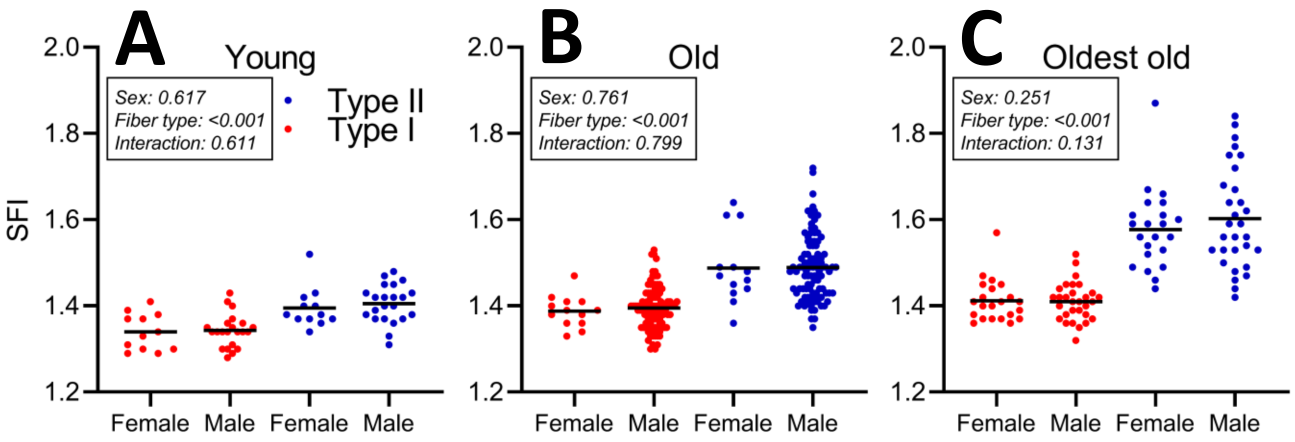
A
Young
Sex: 0.617
Fiber type: <0.001
Interaction: 0.611
Type II
Type I
SFI
2.0
1.8
1.6
1.4
1.2
Female
Male
Female
Male
B
Old
Sex: 0.761
Fiber type: <0.001
Interaction: 0.799
C
Oldest old
Sex: 0.251
Fiber type: <0.001
Interaction: 0.131

## SM 4

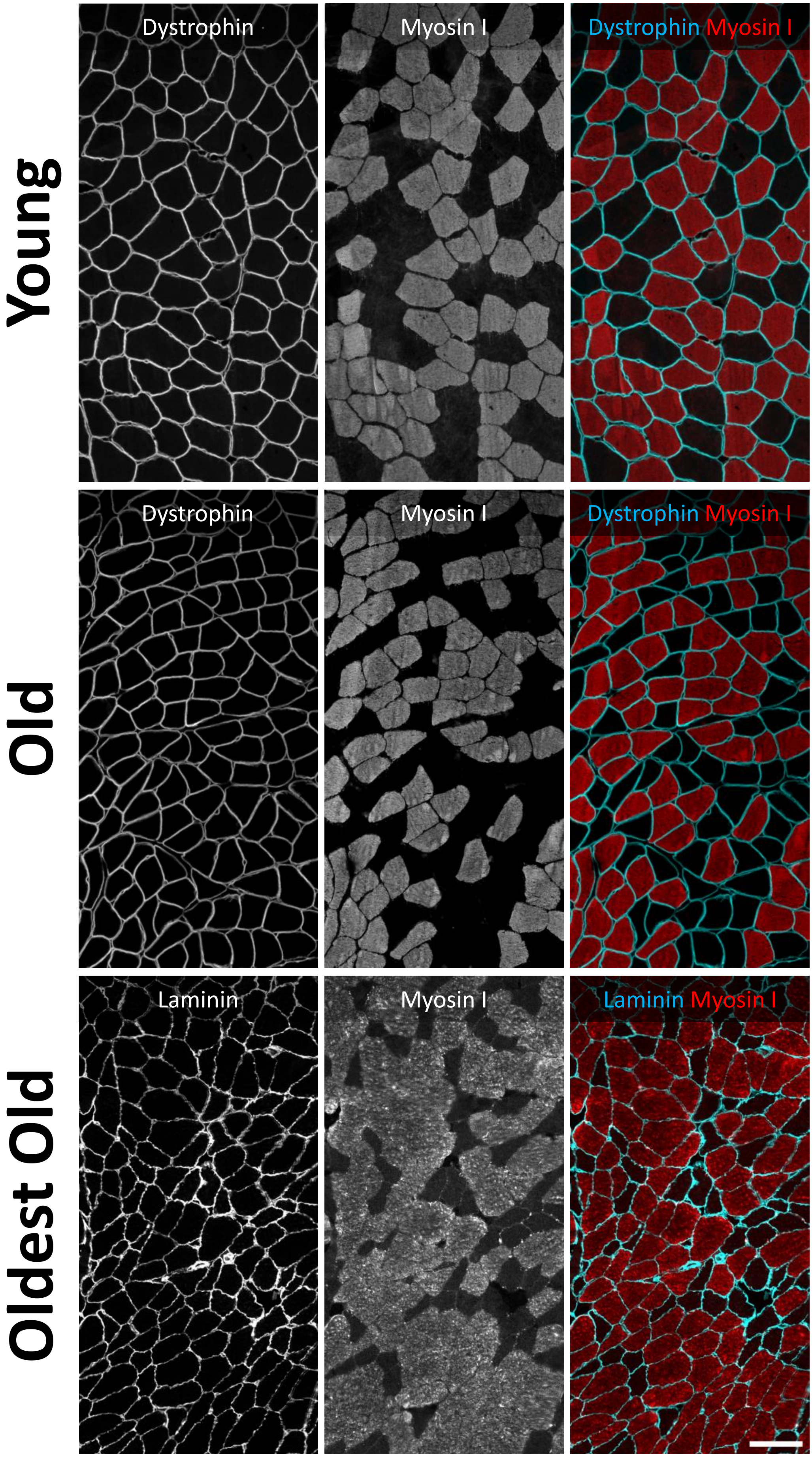
Young
Old
Oldest Old
Dystrophin
Myosin I
Dystrophin Myosin I
Dystrophin
Myosin I
Dystrophin Myosin I
Laminin
Myosin I
Laminin Myosin I

# SM 5

# A

| | | Number of subjects in each SFI increment | | | Number of myofibers in each SFI increment | | |
|---|---|---|---|---|---|---|---|
| | | Young | Old | Oldest Old | Young | Old | Oldest Old |
| Type I | <.1.10 | 1 | 1 | 1 | 1 | 1 | 1 |
| | 1.10-1.20 | 34 | 99 | 49 | 497 | 992 | 316 |
| | 1.20-1.30 | 34 | 111 | 52 | 2859 | 5916 | 2264 |
| | 1.30-1.40 | 34 | 111 | 52 | 2134 | 6041 | 2663 |
| | 1.40-1.50 | 34 | 111 | 52 | 984 | 3576 | 1760 |
| | 1.50-1.60 | 34 | 107 | 51 | 421 | 1845 | 977 |
| | 1.60-1.70 | 30 | 101 | 52 | 144 | 946 | 505 |
| | 1.70-1.80 | 21 | 93 | 45 | 85 | 464 | 290 |
| | 1.80-1.90 | 14 | 76 | 41 | 33 | 253 | 161 |
| | 1.90-2.00 | 7 | 56 | 34 | 10 | 128 | 103 |
| | >2.00 | 16 | 67 | 40 | 35 | 233 | 184 |
| Type II | <.1.10 | 0 | 0 | 1 | 0 | 0 | 1 |
| | 1.10-1.20 | 29 | 79 | 34 | 191 | 410 | 142 |
| | 1.20-1.30 | 34 | 111 | 52 | 1980 | 2924 | 1012 |
| | 1.30-1.40 | 34 | 111 | 52 | 2310 | 4303 | 1705 |
| | 1.40-1.50 | 34 | 111 | 52 | 1326 | 3266 | 1495 |
| | 1.50-1.60 | 34 | 111 | 52 | 673 | 2272 | 1154 |
| | 1.60-1.70 | 33 | 108 | 52 | 306 | 1287 | 774 |
| | 1.70-1.80 | 31 | 104 | 51 | 161 | 912 | 512 |
| | 1.80-1.90 | 27 | 96 | 50 | 75 | 537 | 385 |
| | 1.90-2.00 | 15 | 88 | 51 | 28 | 336 | 238 |
| | >2.00 | 26 | 95 | 51 | 68 | 776 | 757 |

# B

| | | Number of subjects in each CSA increment | | | Number of myofibers in each CSA increment | | |
|---|---|---|---|---|---|---|---|
| | | Young | Old | Oldest Old | Young | Old | Oldest Old |
| Type I | 0-1000 | 10 | 37 | 31 | 73 | 188 | 178 |
| | 1000-2000 | 23 | 78 | 49 | 423 | 1126 | 647 |
| | 2000-3000 | 33 | 99 | 52 | 1256 | 3574 | 1447 |
| | 3000-4000 | 34 | 110 | 51 | 1692 | 4641 | 2015 |
| | 4000-5000 | 34 | 110 | 49 | 1526 | 4421 | 1812 |
| | 5000-6000 | 31 | 109 | 45 | 1068 | 3187 | 1413 |
| | >6000 | 29 | 103 | 41 | 1166 | 3407 | 1734 |
| Type II | 0-1000 | 7 | 52 | 46 | 35 | 923 | 1675 |
| | 1000-2000 | 16 | 92 | 50 | 313 | 2803 | 2707 |
| | 2000-3000 | 27 | 107 | 50 | 898 | 4548 | 1824 |
| | 3000-4000 | 33 | 110 | 48 | 1447 | 4198 | 1048 |
| | 4000-5000 | 33 | 105 | 41 | 1634 | 2418 | 517 |
| | 5000-6000 | 29 | 96 | 27 | 1422 | 1132 | 234 |
| | >6000 | 22 | 66 | 17 | 1371 | 1018 | 187 |

# SM 6

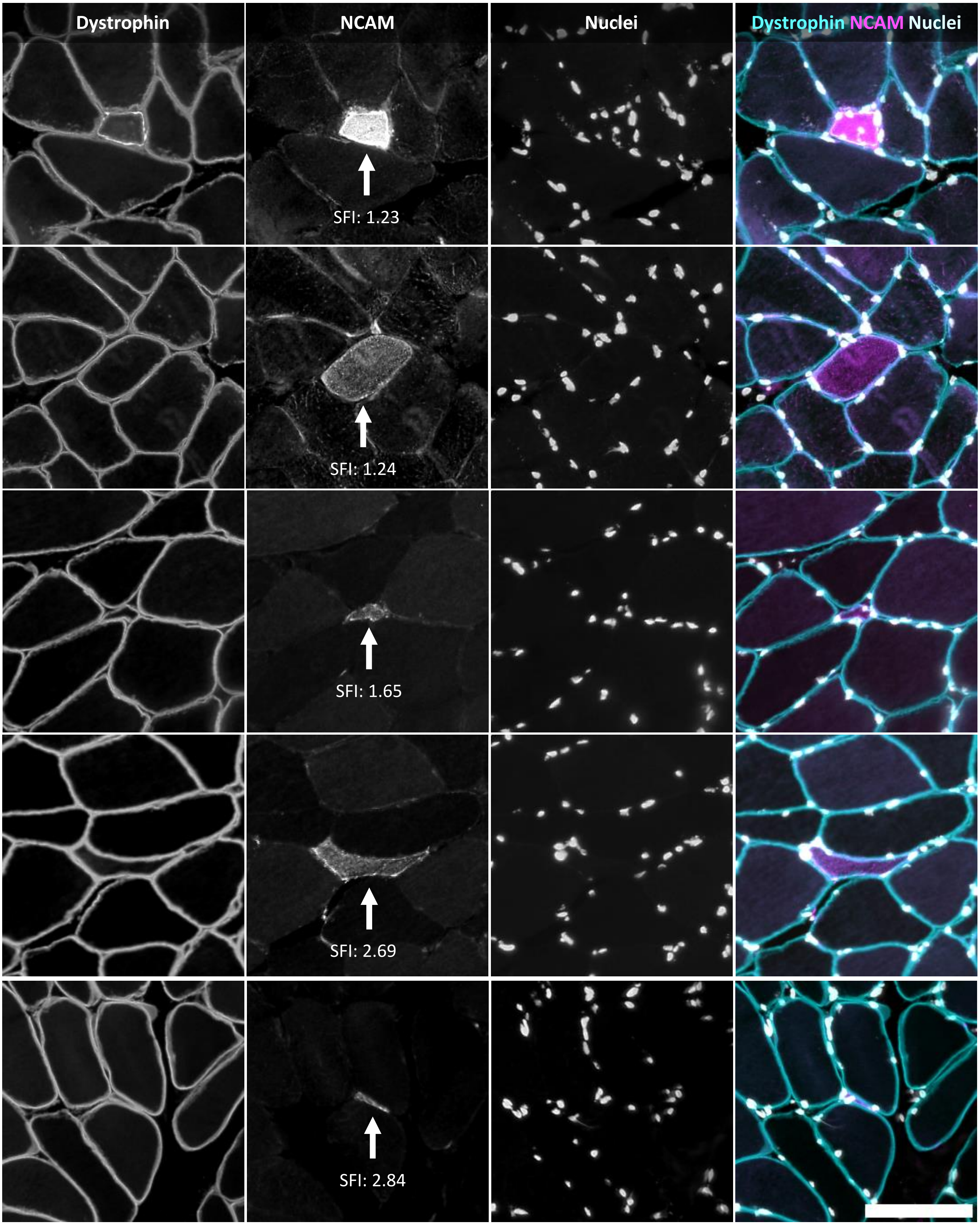
Dystrophin
NCAM
Nuclei
Dystrophin NCAM Nuclei
SFI: 1.23
SFI: 1.24
SFI: 1.65
SFI: 2.69
SFI: 2.84

# SM 7

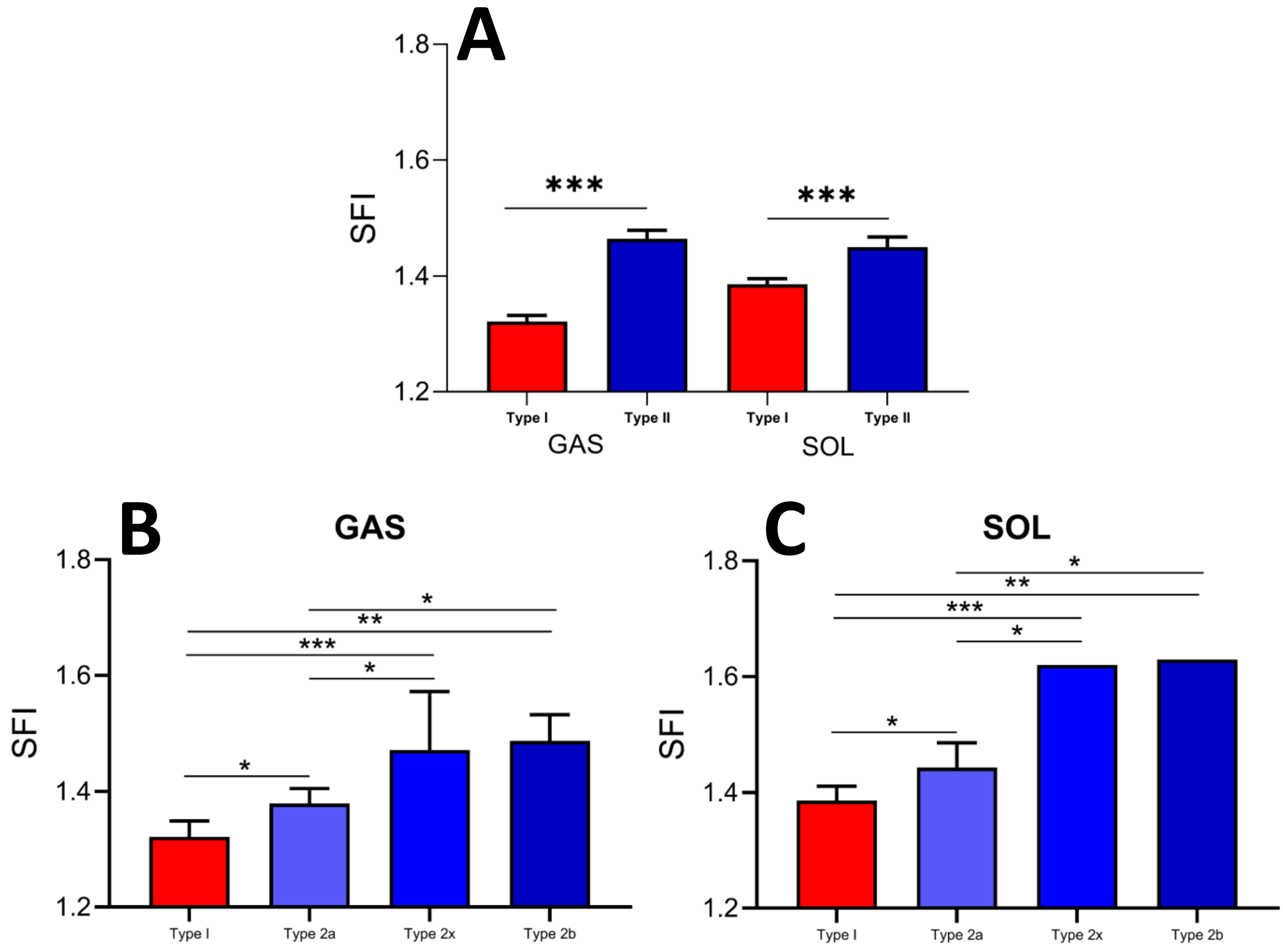

A
1.8
1.6
1.4
1.2
SFI
***
***
Type I
Type II
Type I
Type II
GAS
SOL
B
GAS
SFI
*
**
***
*
*
Type I
Type 2a
Type 2x
Type 2b
C
SOL
SFI
*
**
***
*
*
Type I
Type 2a
Type 2x
Type 2b

# SM 8

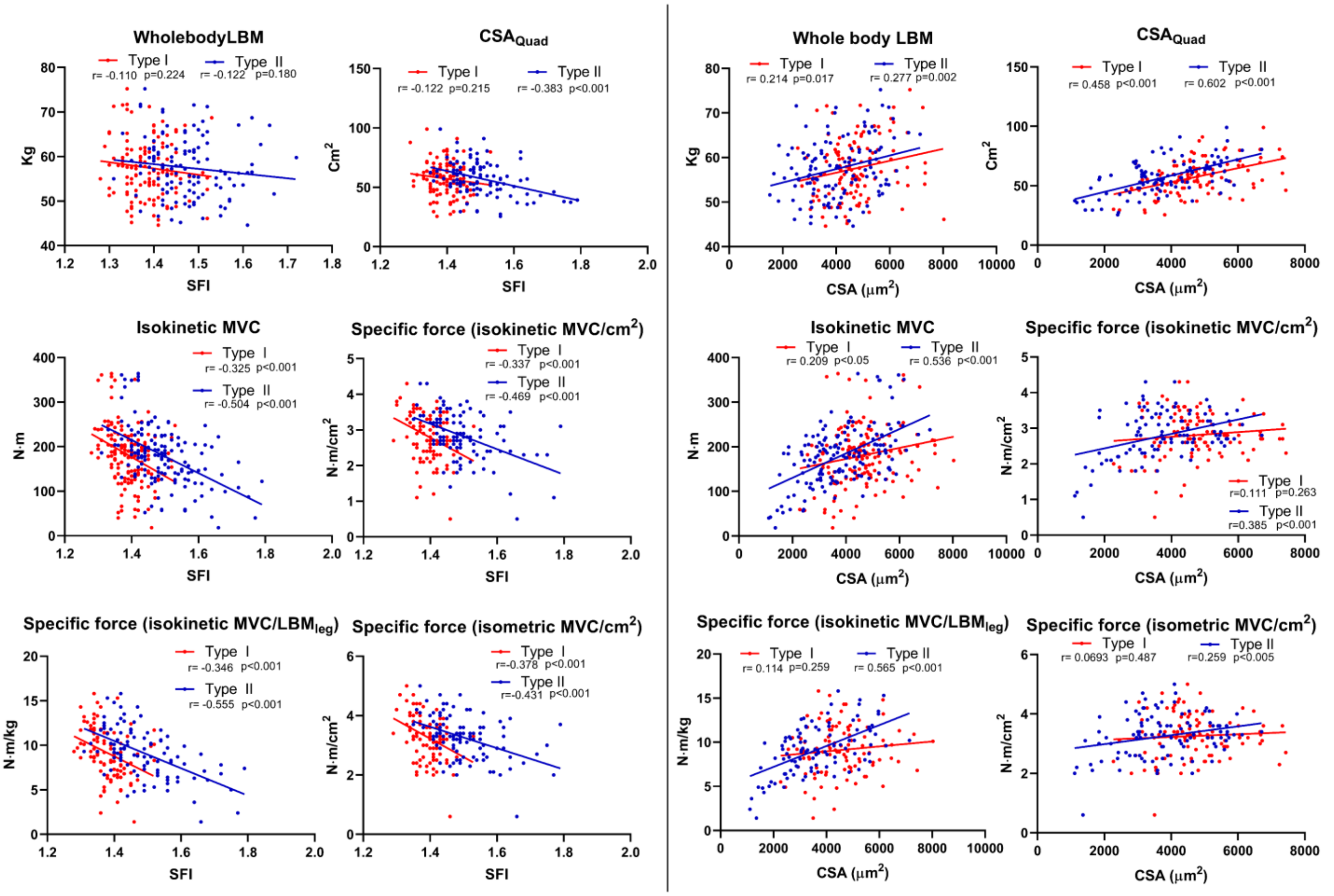

WholebodyLBM
Type I r= -0.110 p=0.224
Type II r= -0.122 p=0.180
Kg
SFI
CSA$_{Quad}$
Type I r= -0.122 p=0.215
Type II r= -0.383 p<0.001
Cm$^2$
SFI
Whole body LBM
Type I r= 0.214 p=0.017
Type II r= 0.277 p=0.002
Kg
CSA (μm$^2$)
CSA$_{Quad}$
Type I r= 0.458 p<0.001
Type II r= 0.602 p<0.001
Cm$^2$
CSA (μm$^2$)
Isokinetic MVC
Type I r= -0.325 p<0.001
Type II r= -0.504 p<0.001
N·m
SFI
Specific force (isokinetic MVC/cm$^2$)
Type I r= -0.337 p<0.001
Type II r= -0.469 p<0.001
N·m/cm$^2$
SFI
Isokinetic MVC
Type I r= 0.209 p<0.05
Type II r= 0.536 p<0.001
N·m
CSA (μm$^2$)
Specific force (isokinetic MVC/cm$^2$)
Type I r=0.111 p=0.263
Type II r=0.385 p<0.001
N·m/cm$^2$
CSA (μm$^2$)
Specific force (isokinetic MVC/LBM$_{leg}$)
Type I r= -0.346 p<0.001
Type II r= -0.555 p<0.001
N·m/kg
SFI
Specific force (isometric MVC/cm$^2$)
Type I r=-0.378 p<0.001
Type II r=-0.431 p<0.001
N·m/cm$^2$
SFI
Specific force (isokinetic MVC/LBM$_{leg}$)
Type I r= 0.114 p=0.259
Type II r= 0.565 p<0.001
N·m/kg
CSA (μm$^2$)
Specific force (isometric MVC/cm$^2$)
Type I r= 0.0693 p=0.487
Type II r=0.259 p<0.005
N·m/cm$^2$
CSA (μm$^2$)

# SM 9

| | LBMleg | | | |
|---|---|---|---|---|
| | *Resulting model: n=108, p<0.0001, R Squared=41%* | | | |
| | **Coefficient** | **SE** | **p-value** | **R Squared** |
| **Intercept** | 32,79 | 8,61 | 0,0002 | |
| **Type I SFI** | -14,23 | 5,97 | 0,0190 | 0,032 |
| **Type II CSA** | 0,002 | 0,0002 | <0,0001 | 0,272 |
| | **Isometric MVC** | | | |
| | *Resulting model: n=150, p<0.0001, R Squared=58%* | | | |
| | **Coefficient** | **SE** | **p-value** | **R Squared** |
| **Intercept** | 579,61 | 69,55 | <0,0001 | |
| **Age** | -2,25 | 0,25 | <0,0001 | 0,229 |
| **Type II SFI** | -187,70 | 53,09 | 0,0005 | 0,036 |
| **Type I CSA** | 0,01 | 0,004 | 0,0013 | 0,031 |
| | **Isometric RFD** | | | |
| | *Resulting model: n=123, p<0.0001, R Squared=35%* | | | |
| | **Coefficient** | **SE** | **p-value** | **R Squared** |
| **Intercept** | 2847,71 | 182,97 | <0,0001 | |
| **Age** | -22,23 | 2,78 | <0,0001 | 0,346 |
| | **Specific force (isometric MVC/LBMleg)** | | | |
| | *Resulting model: n=98, p<0.0001, R Squared=57%* | | | |
| | **Coefficient** | **SE** | **p-value** | **R Squared** |
| **Intercept** | 23,99 | 3,23 | <0,0001 | |
| **Age** | -0,08 | 0,01 | <0,0001 | 0,270 |
| **Type II SFI** | -5,49 | 2,40 | 0,0242 | 0,024 |
| | **Whole body LBM** | | | |
| | *Resulting model: n=123, p=0.0022, R Squared=8%* | | | |
| | **Coefficient** | **SE** | **p-value** | **R Squared** |
| **Intercept** | 51,26 | 2,08 | <0,0001 | |
| **Type II CSA** | 0,002 | 0,0005 | 0,0022 | 0,075 |
| | **CSAquad** | | | |
| | *Resulting model: n=104, p<0.0001, R Squared=55%* | | | |
| | **Coefficient** | **SE** | **p-value** | **R Squared** |
| **Intercept** | 98,32 | 19,64 | <0,0001 | |
| **Age** | -0,26 | 0,08 | 0,0016 | 0,048 |
| **Type II SFI** | -36,30 | 14,00 | 0,0110 | 0,030 |
| **Type I CSA** | 0,004 | 0,001 | 0,0002 | 0,068 |
| **Type II CSA** | 0,003 | 0,001 | 0,0081 | 0,033 |
| | **Isokinetic MVC** | | | |
| | *Resulting model: n=150, p<0.0001, R Squared=57%* | | | |
| | **Coefficient** | **SE** | **p-value** | **R Squared** |
| **Intercept** | 516,30 | 64,08 | <0,0001 | |
| **Age** | -2,04 | 0,23 | <0,0001 | 0,231 |
| **Type II SFI** | -164,82 | 48,84 | 0,0009 | 0,034 |
| **Type I CSA** | 0,01 | 0,003 | 0,0024 | 0,028 |
| | **Specific force (isokinetic MVC/LBMleg)** | | | |
| | *Resulting model: n=99, p<0.0001, R Squared=57%* | | | |
| | **Coefficient** | **SE** | **p-value** | **R Squared** |
| **Intercept** | 22,13 | 3,06 | <0,0001 | |
| **Age** | -0,08 | 0,01 | <0,0001 | 0,264 |
| **Type II SFI** | -5,33 | 2,27 | 0,0209 | 0,025 |
| | **Specific force (isokinetic MVC/CSAquad)** | | | |
| | *Resulting model: n=103, p<0.0001, R Squared=34%* | | | |
| | **Coefficient** | **SE** | **p-value** | **R Squared** |
| **Intercept** | 9,78 | 1,72 | <0,0001 | |
| **Type I SFI** | -3,60 | 1,47 | 0,0164 | 0,040 |
| **Type II SFI** | -1,79 | 0,80 | 0,0273 | 0,034 |
| **Type II CSA** | 0,0002 | 0,00005 | 0,0002 | 0,100 |
| | **Specific force (isometric MVC/CSAquad)** | | | |
| | *Resulting model: n=103, p<0.0001, R Squared=26%* | | | |
| | **Coefficient** | **SE** | **p-value** | **R Squared** |
| **Intercept** | 7,79 | 1,11 | <0,0001 | |
| **Age** | -0,01 | 0,00 | 0,0031 | 0,068 |
| **Type II SFI** | -2,33 | 0,83 | 0,0060 | 0,059 |

# SM 10

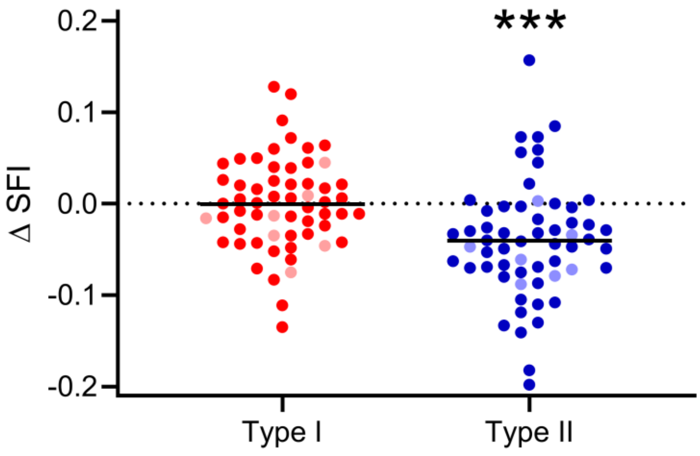
